# Deconstructing biodiversity-ecosystem function relationships: Filtering of macroinvertebrate traits in a large river floodplain

**DOI:** 10.1101/2020.07.15.204008

**Authors:** Natalie K. Rideout, Zacchaeus G. Compson, Wendy A. Monk, Meghann R. Bruce, Mehrdad Hajibabaei, Teresita M. Porter, Michael T.G. Wright, Donald J. Baird

**Affiliations:** Canadian Rivers Institute, Department of Biology, University of New Brunswick, Fredericton, NB, Canada; Environment and Climate Change Canada @ Canadian Rivers Institute, Department of Biology, University of New Brunswick, Fredericton, NB, Canada; Environment and Climate Change Canada @ Canadian Rivers Institute, Faculty of Forestry and Environmental Management, University of New Brunswick, Fredericton, NB, Canada; Canadian Rivers Institute @ University of New Brunswick, Fredericton, NB, Canada; Centre for Biodiversity Genomics and Department of Integrative Biology, University of Guelph, ON, Canada

## Abstract

The Biodiversity-Ecosystem Function hypothesis postulates that higher biodiversity is correlated with ecosystem function by providing a high number of filled niches through species response types and resource use patterns. Through their high spatio-temporal habitat diversity, floodplains are highly productive ecosystems, supporting communities that are naturally resilient and highly diverse. We examined linkages among floodplain wetland habitats, invertebrate communities and their associated traits, and ecosystem function across 60 sites within the floodplain wetlands of the lower Wolastoq | Saint John River, New Brunswick, using structural equation modelling and Threshold Indicator Taxa ANalysis (TITAN2). We identified key environmental filters of invertebrate communities, namely linking increased niche differentiation through historical change, flood pulse dynamics, and macrophyte bed complexity with increased taxa and functional diversity. Examination of traits linked to ecosystem functions revealed that healthy wetlands with higher primary productivity were associated with greater functional evenness and richness, while habitat patches with increased decomposition rates had low functional richness, reflecting highly disturbed habitat. Our results highlight key differences between wetland and riverine ecosystems, relating to how critical functions support healthy wetland habitats by providing increased resilience to disturbance, here associated with differing levels of conservation protection.

## Introduction

In dynamic ecosystems such as floodplains, the ability of communities to support basic ecosystem functions and thus remain resilient to natural and human-induced disturbances is hypothesized to be underpinned by functional diversity—a key component of biodiversity [1]. Functional diversity in turn supports functional redundancy, providing an “insurance policy”, whereby many species exhibit a range of responses to disturbance, ensuring that functions are maintained even as species are lost [2,3]. In a review of 100 studies, Srivastava & Vellend [4] found that 71% reported higher rates of ecosystem function with increased biodiversity. Ecosystem function, often synonymous with ecosystem health, supports an ecosystem’s stability, its ability to maintain energy fluxes (e.g. production and decomposition), and its stocks of energy and biomass [4–8]. In wetlands, healthy functions are vital in linking terrestrial and aquatic food webs through decomposition [9], while aquatic macrophyte and periphyton communities generate energy through primary production [10].

Although the link between biodiversity and ecosystem function has been recognized by ecologists for decades [1], only recently have the mechanisms behind this link been examined, focusing chiefly on functional traits [2]. Traits are defined as the “morphological, physiological or behavioural characteristics of a species that describe a species’ physical characteristics, functional role in an ecosystem, or its ecological niche” [11]. The shift in focus from taxonomy-based biodiversity to traits-based studies is important in that traits allow ecologists to compare across broader scales, where species may be interchangeable, but traits are retained, and account for species that may fill several niches depending on their life stage [12]. Traits-based ecology also encompasses the fact that abiotic variables act as environmental filters primarily for traits, only secondarily filtering for the taxa that hold those traits [13].

From a biomonitoring perspective, it is traits that have an impact on ecosystem functions that are the most critical to conserve [14]; equally important is the maintenance of functional redundancy to account for future disturbances [15]. Ecosystem vulnerability depends on the phylogenetic similarity of groups that provide certain functions (known as effect traits), as environmental filtering for response traits can eradicate entire lineages with similar functional roles [14,16]. In fact, it is response traits—those that influence a species’ ability to colonize and thrive in an environment (and thus, its fitness)—that are subject to natural selection [14]. Trisos, Petchey & Tobias [17], proposed that under high disturbance, communities are dominated by phylogenetically similar species, while communities with low levels of disturbance tend to be more distantly related, reducing competition through niche overlap, resulting in more efficient use of available resources [16]. This theory assumes that selection of species through environmental filtering of response traits will result in communities with similar sets of traits, and therefore lineages, that are more capable of withstanding disturbance [17]. Following this logic, depending on what species and traits dominate within communities, it may be possible to assess the impact of disturbance regimes on biotic communities.

Healthy floodplains support mosaics of habitat patches at different successional stages with varying degrees of connectivity [18,19]. For this reason, they provide an excellent proving ground for traits-based ecological theory, allowing exploration of the linkages between wetland biota and ecosystem function, and the associated role of habitat-based environmental filtering. Despite the rise in traits-based science, taxonomic resolution has imposed limitations [20,21], especially in taxonomically-rich floodplain wetlands [e.g. 22,23], which are less studied compared to in-channel riverine habitats [21]. DNA metabarcoding via high-throughput sequencing provides a powerful tool to characterize community composition in unprecedented detail [24], delivering sufficient taxonomic detail to study how environmental filters, including disturbance, interact with invertebrate traits, and the consequences for maintenance of healthy ecosystem function.

The objective of this study was to determine the drivers of macroinvertebrate community structure and associated wetland ecosystem function. This was done by 1) Quantifying the linkages among environmental filters, macroinvertebrate community structure, and ecosystem function; 2) Determining if there are taxa and trait indicators that can predict habitat change and ecosystem health in floodplain wetlands; and 3) Comparing how taxa and traits respond differently to gradients of function and environmental filtering. In answering these questions, we can examine the link between trait diversity and ecosystem function, and assess how this understanding can help us in biomonitoring for ecosystem health.

## Methods

### Study system

The Wolastoq | Saint John River drains a catchment of 55,110 km^2^ as it flows 673 km from its headwaters in northern Maine and Quebec. Each spring, ice-jams and snow melt generate flood pulses, pushing nutrient and sediment rich water into downstream floodplain wetlands, one of the last semi-intact large-river floodplains in eastern North America. The study area focused on part of this floodplain, Atlantic Canada’s largest freshwater wetland complex, the Grand Lake Meadows and Portobello Creek wetland complex (henceforth abbreviated as the “GLM complex”; Fig 1). These wetlands are a vital reservoir for rare and endangered biodiversity and act as important nursery, flyway, and nesting habitat for many migratory and transitional species, prompting the provincial and federal governments to protect them through the Grand Lake Meadows Protected Natural Area (GLM PNA) and the Portobello Creek National Wildlife Area (PC NWA), respectively. Despite its outstanding conservation value, however, the GLM complex has experienced substantial wetland habitat change in the last few decades, with an altered hydrologic regime, and subsequent sediment and nutrient deposition, into the floodplain due to significant anthropogenic alterations within the watershed [25].

**Fig 1.**
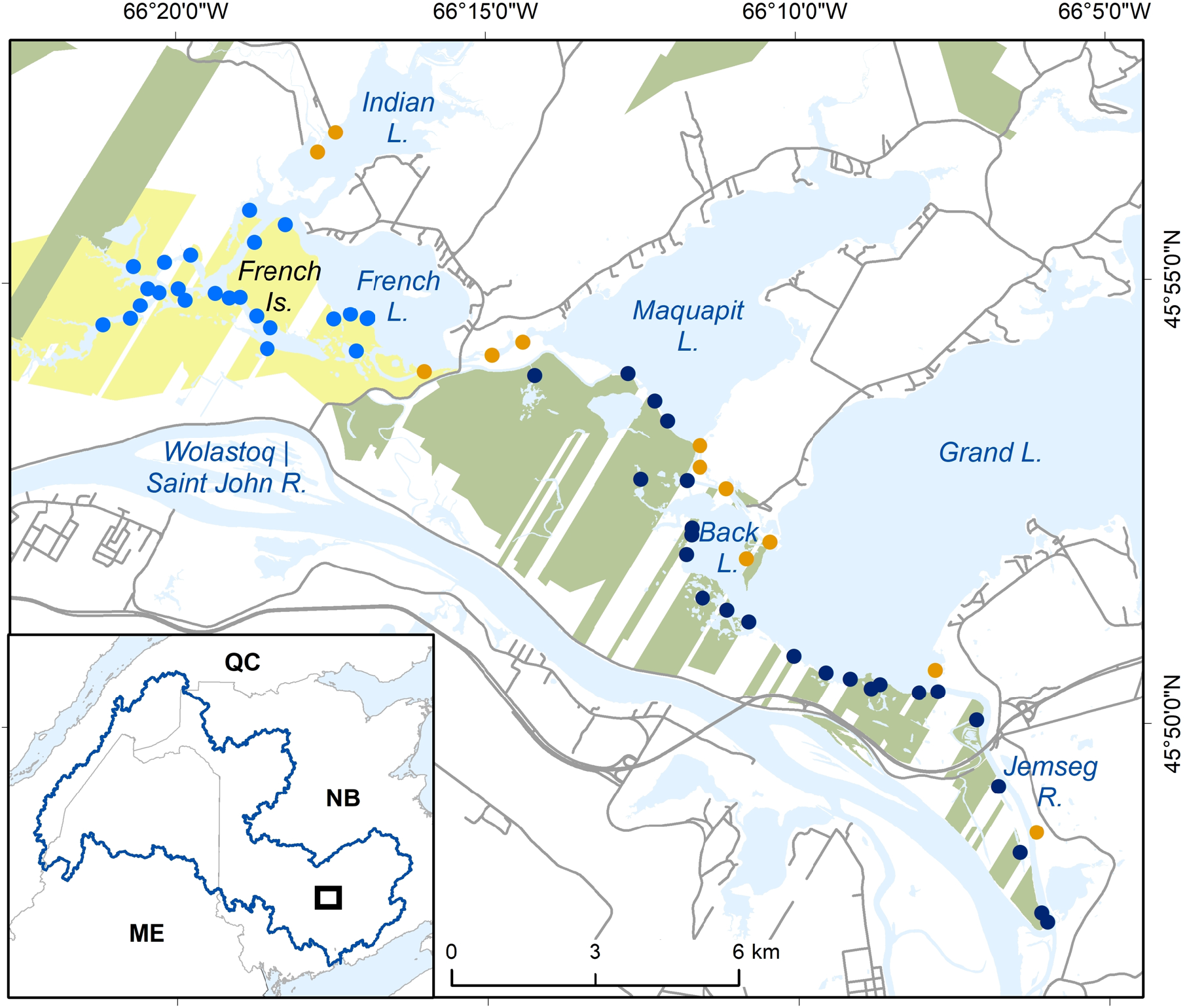
Study sites within the GLM complex relative to different protection strategies. Sites are grouped into Portobello Creek NWA sites (blue), Grand Lake Meadows PNA sites (navy) and sites in areas of no known protection strategy (i.e. “unprotected”) (orange). Major water bodies are labelled and protected areas are coloured: Portobello Creek NWA (yellow) and Grand Lake Meadows PNA (green). Inset map shows location of sampling (black box) in relation to the Wolastoq | Saint John River watershed.

Sixty sites were sampled between June 2017 and August 2017, with sites distributed across the edge of the wetland complex among three levels of protection: 1) unprotected (n = 12), 2) noncontiguous protection (GLM PNA; n = 24; Scientific Protected Natural Area Permit # SCP2016-002 granted to NR by New Brunswick Department of Energy and Resource Devolopment), and 3) contiguous protection (PC NWA; n = 24; National Wildlife Area Permit # NWA3002 granted to NR by the Canadian Wildlife Service) (Fig 1). All sites were characterized as aquatic wetland habitat that extended from the edge of terrestrial high marsh vegetation to open water, containing emergent and submerged macrophytes. At each site, a pair of bamboo poles served to hold sampling equipment and mark the midpoints of each site; all biotic and abiotic samples and surveys were taken within 50 m of these poles.

### Wetland habitat

Part of the supporting habitat data for this work are described in detail in another paper [25], and thus the methods employed are presented in abbreviated form here.

#### Historical change

Total historical change, assessed as the difference in wetland extent between 1951 and 2014 was measured from aerial photos of the study area using ArcGIS (version 10.6.1; [26]). Differences between the two years (1951 and 2014) were calculated for points along the shoreline (n = 2500 transects), from which the entire shoreline was interpolated. Absolute values of total change (in m) for each site were extracted from the resulting interpolated raster file.

#### Abiotic variables

Water and sediment samples were collected following the Canadian Aquatic Biomonitoring Network (CABIN) wetland protocol standard operating procedures [27] and sent to Environment and Climate Change Canada’s National Laboratory for Environmental Testing (NLET) and New Brunswick’s provincial research organization (RPC) to analyse for trace elements, nutrients, organic carbon and physical properties [28,29]. HOBO loggers measured temperature (P/N UA 001 64) and water level (P/N 20-001-04), then summarised as descriptive variables. Hydrologic variables were calculated from water depth data using the Indicators of Hydrologic Alteration software [30].

#### Submerged and emergent macrophyte community

Surveys for emergent and submerged macrophytes were conducted, where submerged macrophytes were defined as aquatic vegetation that were either wholly submerged (e.g. water milfoil) or floating (e.g. pond lilies) and could be rooted or unrooted (e.g. duckweed), while emergent macrophytes were those with roots in the sediment, but most of the plant out of water (e.g. arrowheads). Observers designated the top three dominant species for both emergent and submerged communities at every site and estimated percent cover for each; total macrophyte coverage at each site was estimated and recorded. Representative specimens for all dominant species were collected, along with all submerged macrophyte morphotypes found, for later taxonomic identification.

### Macroinvertebrate community

Benthic macroinvertebrate samples were collected following the CABIN wetland protocol, sweeping through submerged and emergent vegetation for two minutes in order to dislodge invertebrates [27]. Samples were rinsed in the field to remove excess sediment and stored in 1 L jars with 95% ethanol. In the lab, all vegetation was rinsed and removed over a 250 μm sieve, and the ethanol was replaced to ensure that the invertebrate DNA signal was not overwhelmed by macrophyte signal and that the ethanol remained undiluted. Samples were stored at −80ºC until transport to the University of Guelph Centre for Biodiversity Genomics.

Samples were homogenized in sterile blenders and the slurry was subsampled into 50 mL conical tubes. Samples were centrifuged, excess preservative ethanol was removed, and residual ethanol was evaporated at 65C (~ 4-8 hrs). Once dry, the homogenate was subsampled into 2 mL lysing matrix tubes (MP Biomedicals, Solon, Ohio) and further homogenized using a MP FastPrep-24 Classic tissue homogenizer (MP Biomedicals). Samples were lysed overnight with Proteinase K, then extracted using a NucleoSpin Tissue Kit (Machery-Nagel, Duren, Germany) according to the manufacturer’s protocol, eluting with 30 μL molecular grade water. Samples were extracted in batches of 12-18 with a negative control (no sample added) for each batch.

Two COI fragments were amplified using the primer sets BR5 (B 5’ CCIGAYATRGCITTYCCICG, ArR5 5’ GTRATIGCICCIGCIARIACIGG) and F230R (LCO1490 5’ GGTCAACAAATCATAAAGATATTGG, 230_R 5’ CTTATRTTRTTTATICGIGGRAAIGC) [24,31–33]. A two-step PCR protocol was used where regular PCR primers (above) were used in the first step. Reactions contained 2μL DNA template, 17.5μL molecular biology grade water, 2.5μL 10× reaction buffer, 1μL MgCl_2_ (50mM), 0.5μL dNTPs mix (10mM), 0.5μL forward primer (10mM), 0.5μL reverse primer (10mM), and 0.5μL Platinum Taq polymerase (Invitrogen). Thermocycler conditions were as follows: 95°C for 5min, followed by a total of 35 cycles of 94°C for 40s, 46°C for 1min, and 72°C for 30s, and a final extension at 72°C for 5min, and hold at 4°C. Two μL of the first PCR products were used for a second PCR with Illlumina-tailed primers. A negative control was included for each batch of PCR, which was carried through each of the two PCR steps. Amplification success was confirmed visually using a 1.5% agarose gel. Amplicons were purified using a MinElute DNA purification system (Qiagen) and quantified using a Quant-iT (Invitrogen, Waltham Massachusetts, United States) PicoGreen dsDNA assay on a TBS-380 Mini-Fluorometer (Turner Biosystems Sunnyvale California, United States). All samples were normalized to the same concentration, and the two amplified fragments were pooled for each sample prior to indexing using the Nextera XT Index Kit (Illumina, SanDiego, California FC-131-1002). Indexed samples were pooled into one tube, purified through magnetic bead purification, and quantified using the PicoGreen dsDNA assay. Average fragment length was determined on an Agilent Bioanalyzer 2100 (Santa Clara, California, United States) before sequencing the library on an Illumina MiSeq using the V3 sequencing chemistry kit (2 × 300, MS-10203003). A 10% spike-in of PhiX was used as a control.

Raw paired-end metabarcode reads were processed using the SCVUC v2 bioinformatic pipeline available from https://github.com/Hajibabaei-Lab/SCVUC_COI_metabarcode_pipeline. Briefly, reads were paired using SeqPrep (St. John J. SeqPrep. Downloaded 2016. Available: https://github.com/jstjohn/SeqPrep/releases) setting a minimum Phred quality score cutoff of 20 and requiring a minimum overlap of 25 bp. The forward and reverse primers were trimmed using CutAdapt v1.13 by trimming the 5’ primer region, requiring a minimum length of at least 150 bp after trimming, setting a minimum read quality of Phred 20 at the ends of the sequences, and allowing a maximum of 3 N’s [34]. Forward primer trimmed reads were then processed to trim the 3’ primer region with the same settings as above. Unique clusters were obtained using VSEARCH v2.50 [35] while tracking the number of reads per cluster. Unique clusters were denoised using USEARCH v10.0.240 with the unoise3 algorithm [36]. The denoising step corrects putative sequence errors, removes putative chimeric sequences, and removes rare sequences, in this case, clusters with only one or two sequences (singletons and doubletons). An ESV × sample table containing read counts was generated in VSEARCH. Denoised reads were taxonomically assigned using the Ribosomal Database Project classifier v2.12 that uses a naive Bayesian method (version 2.12; [37]) with a custom COI reference sequence database [38]. Assignments were then filtered for greater than 99% confidence for correct assignments at the genus level. Data generated from DNA-metabarcoding were treated as presence/absence information [39].

Ecologically relevant traits were chosen to describe the niches and response patterns within the wetland ecosystem; chosen traits were limited to those with information that was feasibly accessible for all taxa. Trait information for 13 traits and 67 modalities (Table 1) were assigned at the genus level primarily from the USGS Database of Lotic Invertebrate Traits for North America [20]. Gaps were filled in first from the European *freshwaterwater.info* database [27], and then from the literature; lastly, any remaining gaps, predominantly amongst zooplankton taxa, were completed at the family level. Traits were assigned as either “effect” (i.e. having a direct impact on ecosystem functions or properties) or “response” (i.e. influencing an organism’s ability to persist under environmental changes) [14] according to their main assumed role in the community.

**Table 1.**
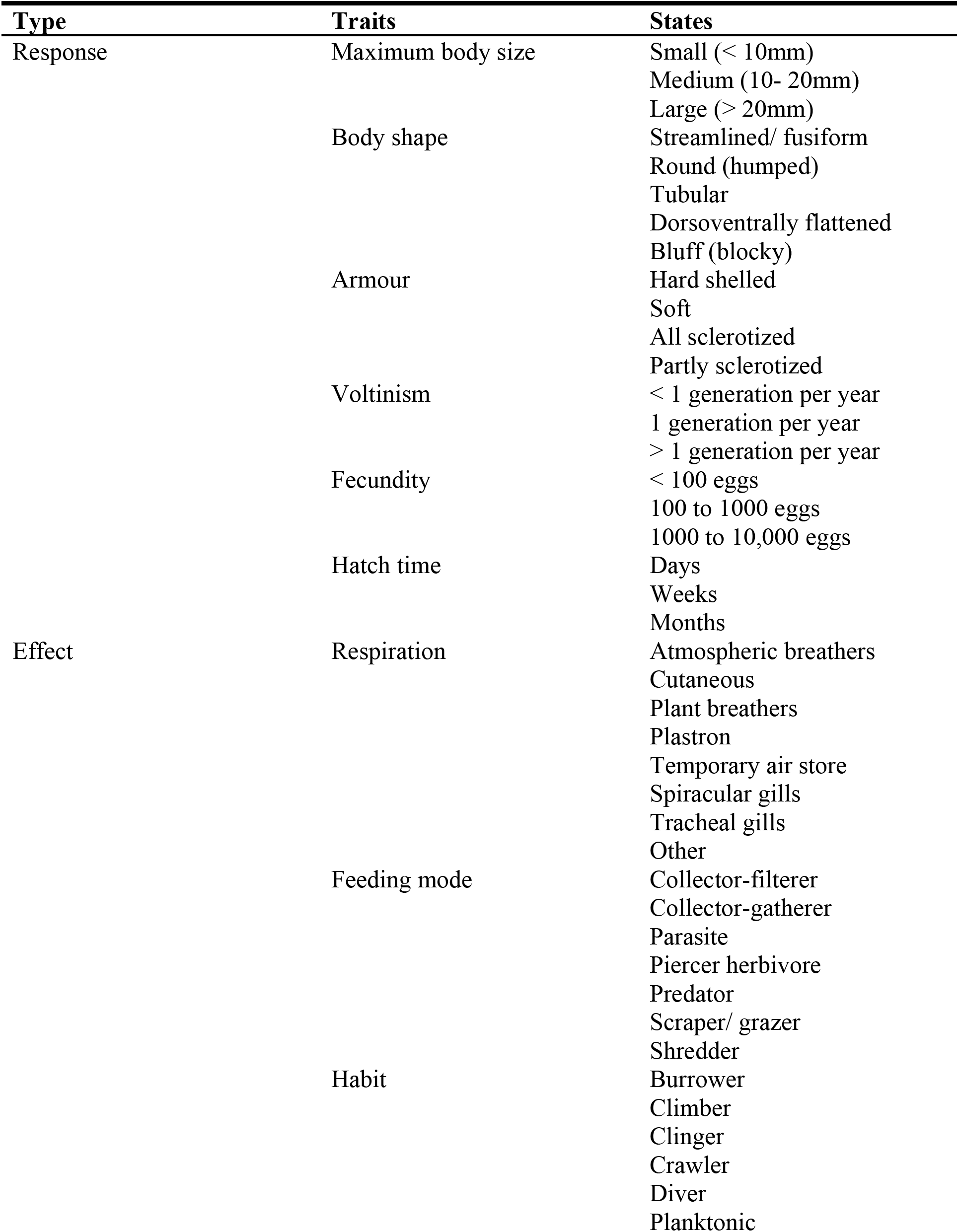
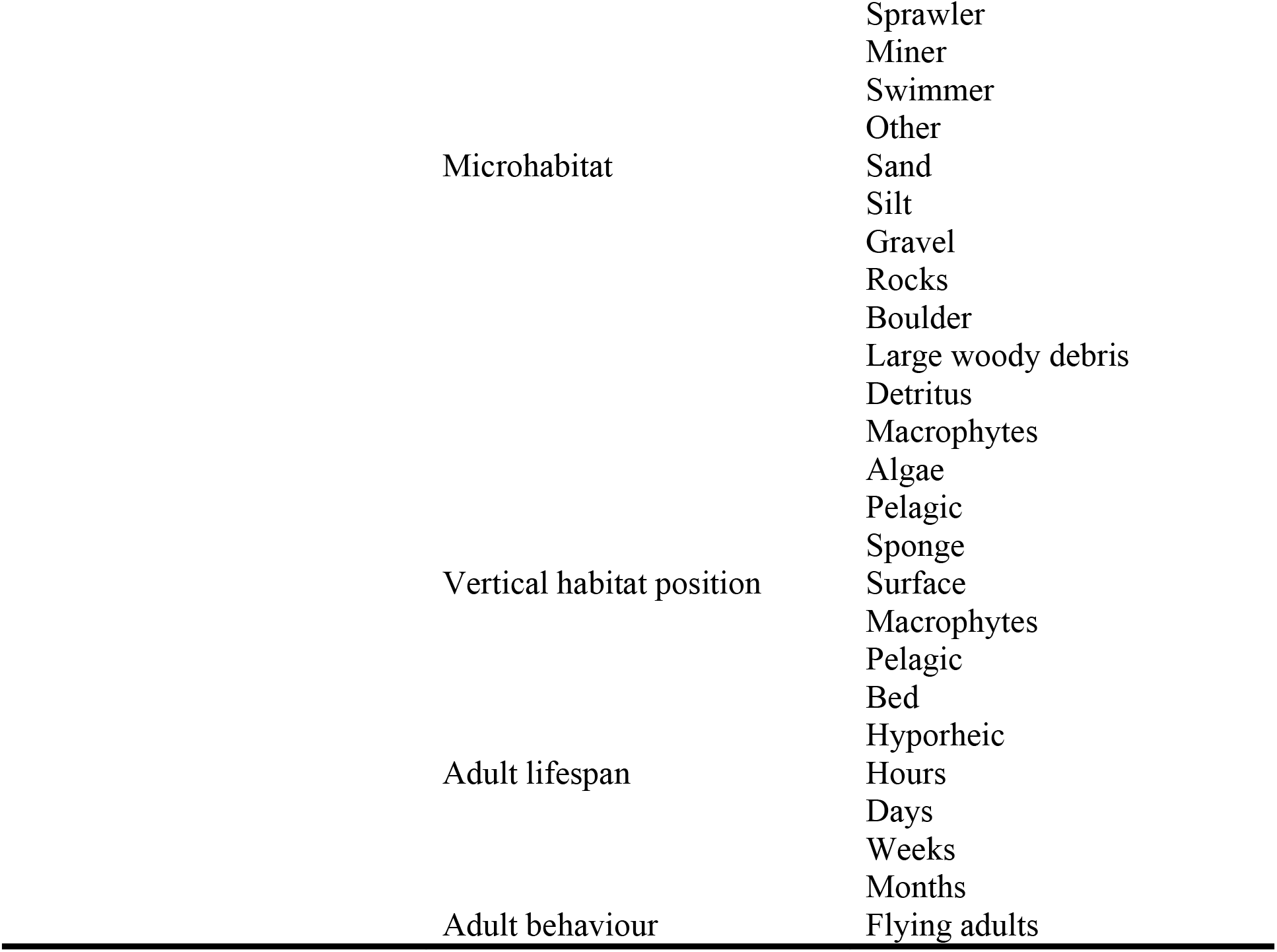
Summary of macroinvertebrate traits and their states that were included in analyses. All taxa were assigned either a 1 or 0 for each state, with multiple states possible for each trait.

### Ecosystem function measures

Primary production was assessed by measuring chlorophyll-*a* levels from unglazed tiles (n = 3 per site) that accumulated algae in the field for 21 days. Each 4.7 × 4.7 cm tile was scraped into de-ionized water, measured onto filter papers, and boiled for 7 minutes at 80ºC to extract chlorophyll-*a*, which was then measured using a Turner Designs Trilogy fluorometer (accuracy μg/L).

Decomposition was assayed with litter packs deployed between June 19 - 23 and incubated for exactly 21 days. Briefly, silver maple (*Acer saccharinum* L. 1753) leaf litter (3.000 ± 0.010 g) was placed into 12.7 × 12.7 cm packs (n = 3 per site) made from 10 mm plastic chicken wire, allowing for the inclusion invertebrates but exclusion of most fish. An extra set of leaf packs (n = 5) was made to account for handling loss and to determine the original percentage of organic matter present in the silver maple leaves. Harvested leaf packs were frozen until they were processed using a modification of the Benfield [41] method. In short, packs were rinsed for excess sediment over a 250 μm sieve and invertebrates picked to reduce errors in weight measurements, then dried for 48 hours at 65ºC and weighed to 0.001 g. Excess suspended sediment in the water column led to many packs having excessively high weights after field collection. To account for this, the organic content of the leaf packs was burned off at 500ºC for 2 hours and ash free dry mass (AFDM) was calculated according to standard methods [41].

### Statistical analysis

To examine linkages among environmental drivers and disturbance, macroinvertebrate community structure, and ecosystem function, structural equation models (SEM) were created in IBM SPSS Amos (version 25.0.0; [42]). SEM is a powerful statistical technique that infers causation between correlative variables by fitting the data to an *a priori* constructed model, while taking all other variables into consideration; it compares the hypothesized model to a random null model to assess model fit [43,44]. The maximum likelihood *χ*^2^ value and its associated *p*-value were examined, where a non-significant result indicated that the hypothesized model was not significantly different from the data confronting the model. The root mean square error approximation (RMSEA) and the goodness of fit index (GFI) were also examined as additional measures of model fit. Since the models were made *a priori* using our best knowledge about the ecosystem, and model fits (as indicated by *p*-values > 0.05, low *χ*^2^ values, and GFI values close to 1) were satisfactory, no further changes were made to the models. Additionally, as this was an exploratory analysis, it was beneficial to examine all pathways, whether or not they were statistically significant in predicting ecosystem responses, as non-significant linkages still provide valuable information, especially for wetland systems, which are understudied compared to their in-channel counterparts.

In order to compare if invertebrate community assemblage or aspects of its functional diversity— richness and evenness—better explained variation in the GLM complex, three single biotic metric models were made. Akaike Information Criterion (AIC) values between the three single biotic metric models were compared to determine the best fitting model and compare which biotic index had the most explanatory power in the ecosystem. SEM is a path-based analysis that considers the linkages between all other variables in its final determination of results; thus, in terms of power, only models with the same number of paths can be compared. An additional structural equation model was made that incorporated the three aspects of functional diversity outlined by Villéger *et al.* [45]—richness, evenness and divergence—in a single model. In total, we tested four SEMs.

All abiotic variables entering the model were reduced for high correlations at a threshold of 0.7 Pearson correlation coefficient and condensed for analysis with Principal Components Analysis (PCA) using the *FactoMineR* package (version 2.3; [46]) in R (version 3.5.3; [47]) (Table 2); three axes were chosen based on assessment of the scree plot, explaining 43.6% of the variation of abiotic variables among sites. Axes were labelled according to their strongest associated variables: PC1 (19.9%) was classified as *“Nutrients”*, PC2, (14.5%) was labelled “*Temperature and Metals*”, and PC3 (8.3%) was denoted as “*Hydrology”.* Emergent and submerged macrophyte species were combined and condensed for analysis using a Principal Coordinates Analysis (PCoA) in *vegan* (version 2.5-4; [48]) in R (version 3.5.3; [47]); two axes were chosen, encompassing 16.6% of the total variation. Macroinvertebrate taxa were also condensed to a single axis using a PCoA, representing 11.8% of the community’s variation. Using the *FD* package (version 1.0-12; [48,49]) in R (version 3.5.3; [47]), the following multidimensional trait diversity metrics were calculated: functional richness (defined by the total trait space), functional evenness (i.e. the evenness in the distribution of traits and their relative prevalence in trait space) and functional divergence (which represents how prevalence is distributed within the trait space, relative to the centroid) [45].

**Table 2.**
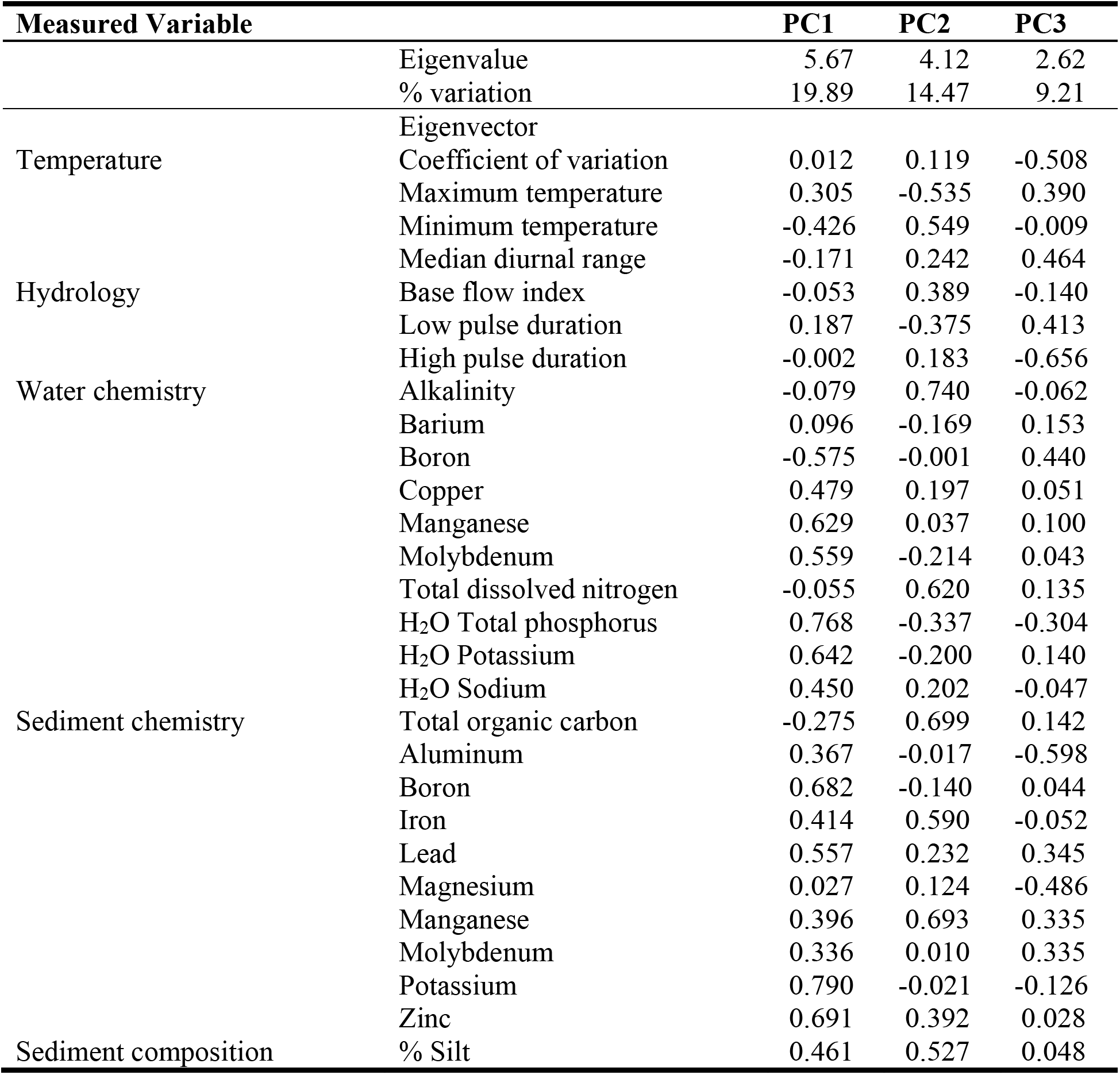
Variable reduction with Principal Component Analysis for abiotic input in structural equation models. Based on the patterns with the strongest associating variables, PC1 was classified as “*Nutrients”* PC2 as “*Temperature and Metals*” and PC3 as “*Hydrology*” for labels in the structural equation models.

Select pathways (i.e., those that drove the differences in the response variables or community dynamics among sites) from the SEMs were examined more closely by breaking composite PCoA and PCA axes into their component variables and then using a Distance-based linear model (DistLM) in the PERMANOVA+ extension for PRIMER (version 7; [51,52]) to establish which variables contributed most to linkages between abiotic and biotic or functional components of the ecosystem. Dissimilarity matrices were first calculated using Euclidean distance for abiotic variables and functional diversity metrics (to maintain congruency with how functional diversity metrics were calculated within the *FD* package) and Bray-Curtis for the biotic community. DistLM uses the dissimilarity matrices from the multivariate predictor and response datasets to determine significant predictor variables using stepwise sequential tests (total permutations = 9999).

To examine trait and taxa indicators of wetland health, namely decomposition and nutrient gradients, we used Threshold Indicator Taxa ANalysis (*TITAN2* version 2.1; [53]) in R (version 3.5.3; [47]). TITAN2 analysis assesses whether species are “pure” (> 95% have the same response direction for 999 bootstrapped runs) and “reliable” (> 95% of bootstrapped runs are significantly different than null at *p* < 0.05) indicators of the gradient, and classifies them as either positive or negative responders, as well as identifying their optimal range along the gradient [54]. This analysis allows assessment of whether ecosystem function and environmental filters act differently on traits, and taxa that express those traits.

## Results

### DNA metabarcoding of wetland macroinvertebrates

The GLM complex is rich in aquatic invertebrate life with DNA metabarcoding identifying a total of 157 genera from 86 families across the wetlands; of those, 120 genera were from within Class Insecta. Distribution in the study area varied, with site richness ranging from 3 to 63 taxa per site (mean = 30.9, SD = 11.93). Two genera, *Amnicola* (a freshwater snail from the family Amnicolidae) and *Sida* (a water flea from the suborder Cladocera) were present at all sampled sites. Individual taxa were found at a mean of 11.81 (SD=14.00) sites, with 77 taxa found at less than 10% of sampled sites. A total of 20 unique zooplankton and 30 Chironomidae genera were identified throughout the study area, a detection notable in that these taxa are generally difficult to identify from morphology alone.

### Relationships between environmental drivers, biotic components and ecosystem function: SEMs

To gain insights into the factors that influenced this wetland diversity, four structural equation models (SEMs) were used to examine biotic components, the physical habitat and disturbance, and ecosystem function within the wetland complex (Figs 2–5; see Table 3 for associated significant pathways for all SEM models; all pathways, including statistically non-significant ones, are provided in the supplementary information (S1)). All four models revealed that historical change was associated with changes to *Hydrology* (PC3), with a shift to decreased low pulse duration and diurnal temperature range, and increased temperature coefficient of variation, high pulse duration and total organic carbon content of the sediment (Table 3). Also common to the models was an association between *Metals and Temperature* (PC2) and measured chlorophyll-*a* for algae tiles, a proxy measurement of primary production (Table 3). Negative principal component scores, associated with higher maximum temperatures and low levels of metals, led to increased chlorophyll-*a* at the site (Table 3).

**Table 3.** **Significant pathways from structural equation models, corresponding to Figs 2–5.** For statistically non-significant pathways, please refer to supplementary material (S1 Table).

| Model | Predictor | Response | $r$ | $p^a$ |
| --- | --- | --- | --- | --- |
| Invertebrate community | Historical change | Hydrology | -0.418 | *** |
|  | Metals + temperature | Dominant macrophytes | 0.252 | 0.037 |
|  | Nutrients | BMI community | -0.324 | *** |
|  | Diverse macrophytes | BMI community | 0.397 | 0.002 |
|  | Dominant macrophytes | BMI community | -0.315 | *** |
|  | Metals + temperature | Chlorophyll a | 0.277 | 0.026 |
|  | BMI community | Decomposition | 0.459 | *** |
| Functional richness | Historical change | Hydrology | -0.418 | *** |
|  | Metals + temperature | Dominant macrophytes | 0.252 | 0.037 |
|  | Nutrients | Functional richness | 0.274 | 0.010 |
|  | Diverse macrophytes | Functional richness | -0.335 | 0.002 |
|  | Metals + temperature | Chlorophyll a | -0.273 | 0.024 |
|  | Functional richness | Decomposition | -0.348 | 0.006 |
| Functional evenness | Historical change | Hydrology | -0.418 | *** |
|  | Metals + temperature | Dominant macrophytes | 0.252 | 0.037 |
|  | Nutrients | Functional evenness | -0.275 | 0.021 |
|  | Diverse macrophytes | Functional evenness | 0.237 | 0.045 |
|  | Metals + temperature | Chlorophyll a | -0.282 | 0.020 |
|  | Nutrients | Chlorophyll a | 0.329 | 0.009 |
|  | Nutrients | Decomposition | -0.262 | 0.035 |
| Functional diversity metrics | Historical change | Hydrology | -0.418 | *** |
|  | Metals + temperature | Dominant macrophytes | 0.252 | 0.037 |
|  | Nutrients | Functional evenness | -0.312 | 0.006 |
|  | Diverse macrophytes | Functional evenness | 0.241 | 0.030 |
|  | Dominant macrophytes | Functional evenness | -0.276 | 0.023 |
|  | Dominant macrophytes | Functional divergence | -0.273 | 0.038 |
|  | Hydrology | Functional richness | -0.274 | 0.017 |
|  | Diverse macrophytes | Functional richness | -0.270 | 0.013 |
|  | Functional divergence | Functional evenness | -0.314 | 0.007 |
|  | Functional evenness | Functional richness | -0.274 | 0.018 |
|  | Functional richness | Decomposition | -0.311 | 0.022 |
|  | Metals + temperature | Chlorophyll a | -0.289 | 0.013 |
|  | Nutrients | Chlorophyll a | 0.301 | 0.017 |
|  | Functional richness | Chlorophyll-a | 0.294 | 0.038 |
|  | Functional evenness | Chlorophyll a | 0.314 | 0.025 |

$r$ values are the standardized regression weights and indicate the strength and direction of the relationship.
<sup>a</sup> $p$ values listed as \*\*\* are significant at $p < 0.001$ .

**Fig 2.**
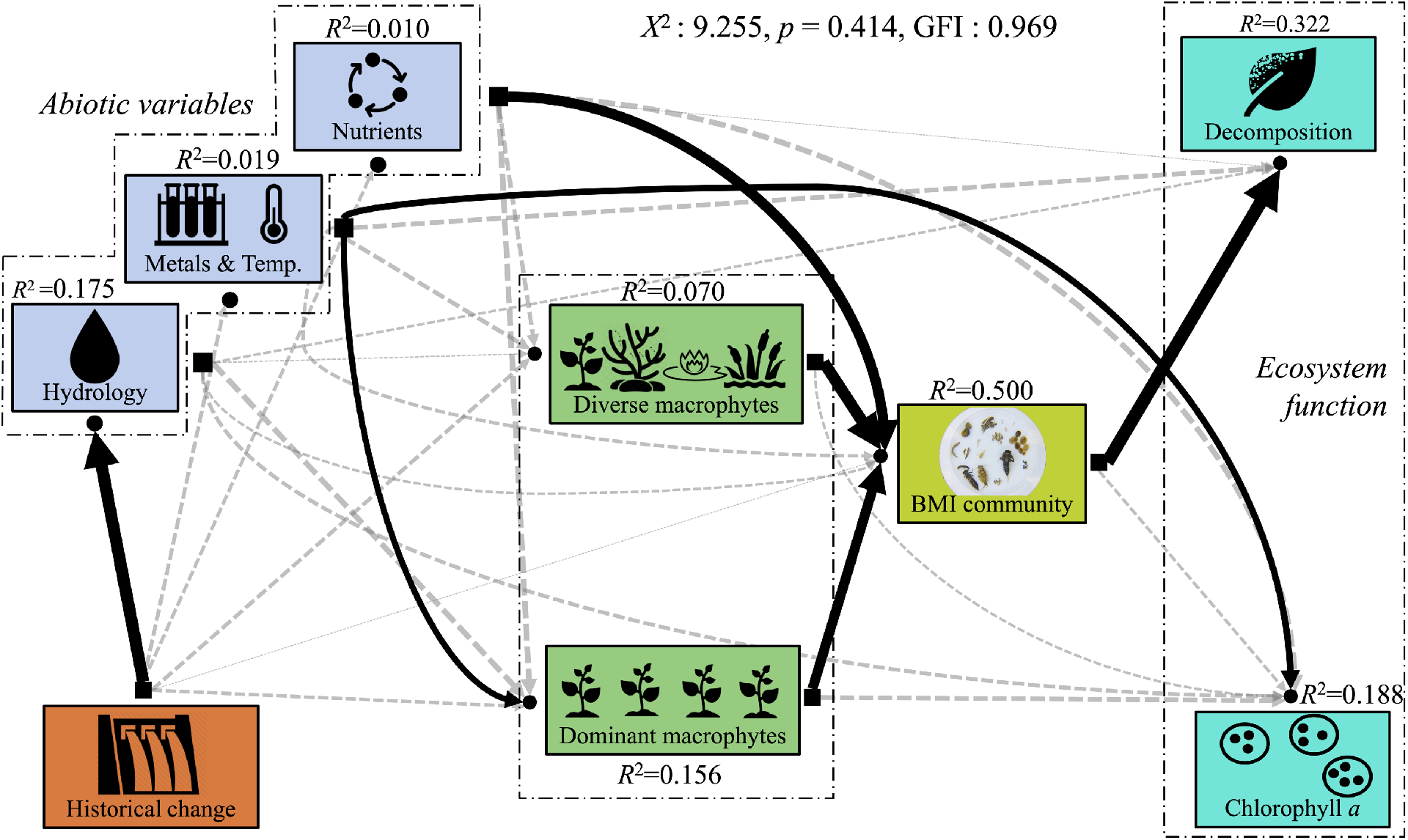
Structural equation model examining the links among the physical habitat, invertebrate community assemblage, and ecosystem function. Black lines represent significant pathways at *p* < 0.05, while statistically non-significant pathways in the model are depicted as dotted, gray lines. The weight of the line represents the strength of the relationship (standardized regression weights, *r*), with thicker lines having a larger effect on explaining the variation of the response variable. The variation of the response variables that are explained by the input variables is shown above each component by its *R*^2^ value.

Macrophytes clearly separated on two principal coordinate axes. PCo1 was associated with species which were common, but when present at a site tended to be associated with higher richness (e.g., *Brasenia schreberi* J.F. Gmel*, Pontederia chordata* L.*, Nuphar lutea* L.(Sm.) and as such is labelled as *Diverse Macrophytes* in the models. PCo2 was associated with emergent and submerged species that, when present, tended to dominate the site with high percent cover (e.g., *Myriophyllum heterophyllum* Michx., *Equisetum fluviatile* L.) and so is denoted as *Dominant Macrophytes.* In all models, the *Metals and Temperature* axis was significant in driving the dynamics of macrophytes that were dominant forming (*Dominant Macrophytes*), with scores indicating higher concentrations of metals associated with the species. DistLM analysis (Table 4) further revealed that the most important abiotic variables for driving the spatial changes in composition of macrophyte communities were: total dissolved nitrogen, alkalinity, copper and molybdenum; total explained variation in both the SEM and the DistLM was low, however, explaining 22.6% and 16.83% of the total variation in the models, respectively.

**Table 4.** DistLM results from selected SEM pathways showing variables driving the differences in the response variable or community dynamics among sites.

|  |  | *r* | *p* |
| --- | --- | --- | --- |
| **Abiotic → Macrophyte community**<br>R<sup>2</sup>=0.1683 | H<sub>2</sub>O Total dissolved nitrogen | 0.0441 | 0.0005 |
|  | H<sub>2</sub>O Alkalinity | 0.0431 | 0.0004 |
|  | H<sub>2</sub>O Copper | 0.0427 | 0.0006 |
|  | SED Molybdenum | 0.0384 | 0.0008 |
| **Abiotic → Invertebrate community**<br>R<sup>2</sup>=0.4749 | H<sub>2</sub>O Manganese | 0.0855 | 0.001 |
|  | H<sub>2</sub>O Alkalinity | 0.0604 | 0.007 |
|  | H<sub>2</sub>O Total Phosphorus | 0.0476 | 0.0029 |
|  | H<sub>2</sub>O Boron | 0.0382 | 0.0036 |
| **Abiotic → Functional richness**<br>R<sup>2</sup>=0.4768 | H<sub>2</sub>O Barium | 0.3081 | 0.0002 |
|  | H<sub>2</sub>O Molybdenum | 0.0330 | 0.0921 |
|  | H<sub>2</sub>O Alkalinity | 0.0555 | 0.0448 |
|  | H<sub>2</sub>O Total Phosphorus | 0.0385 | 0.0533 |
|  | H<sub>2</sub>O Aluminum | 0.0418 | 0.0473 |
| **Abiotic → Functional evenness**<br>R<sup>2</sup>=0.1750 | H<sub>2</sub>O Total dissolved nitrogen | 0.1298 | 0.0039 |
|  | SED Manganese | 0.0452 | 0.082 |
| **FD metrics → Decomposition**<br>R<sup>2</sup>=0.2016 | Functional richness | 0.2016 | 0.0012 |

R<sup>2</sup> represents the total variation explained in the response dataset by the predictor dataset; *r* represents the proportion of variance explained by the individual predictor variable.

The invertebrate assemblage model (Fig 2; Table 3) had the best overall fit (AIC = 81.225) and had the most explanatory power of the single biotic metric models, despite only including one PCoA axis. The model explained 50% of the variation in the invertebrate community, which was driven by higher levels of macro- and micro-nutrients (i.e., those associated with PC1 (*Nutrients*), namely increased total dissolved nitrogen, total phosphorus, copper, molybdenum, and potassium), as well as macrophyte community structure. Both aspects of plant communities (i.e., richness and habitat formation) were associated with changes in invertebrate dynamics, leading to increased diversity (high PCoA scores, including rarer species), with *Diverse Macrophytes* having the strongest pathway and thus explaining most of the total variation. DistLM results (Table 4) indicated that within the abiotic variables, manganese, alkalinity, total phosphorus and boron significantly predicted species differences among wetland sites, explaining 47.5% of the total variation.

In the functional richness model (AIC = 84.780; Fig 3; Table 3), richness was influenced by hydrology, nutrients, and presence of macrophytes, with the model explaining 33.9% of the total variation. As with the invertebrate assemblage model, principal component scores associated with higher levels of nutrients and were associated with increased functional richness. Higher levels of functional richness were associated with decreased low pulse duration and diurnal temperature range, and increased temperature coefficient of variation, high pulse duration, and total organic carbon content of the sediment, suggesting a positive, indirect pathway between historical change and hydrology on functional richness. Macrophytes associated with diverse plant communities also increased trait richness. DistLM results (Table 4) indicated that the most important abiotic variables in driving changes in functional richness were manganese concentration, alkalinity, total phosphorus and boron concentrations (*R*^2^ = 0.48).

**Fig 3.**
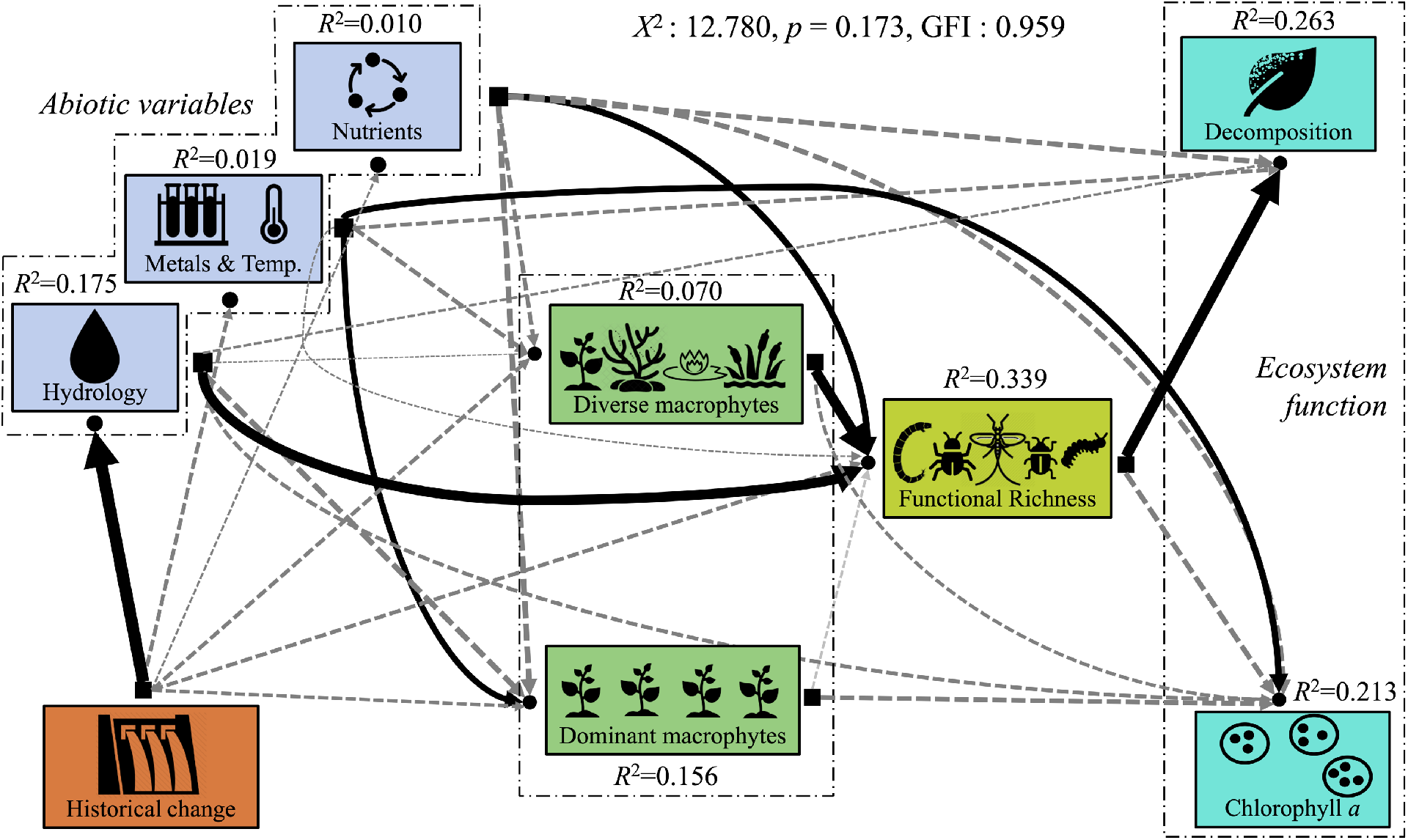
Structural equation model examining the links among the physical habitat, functional richness, and ecosystem function. Black lines represent significant pathways at *p* < 0.05, while statistically non-significant pathways that were included in the model are depicted as dotted, gray lines. The weight of the line represents the strength of the relationship (standardized regression weights, *r*), with thicker lines having a larger effect on explaining the variation of the response variable. The variation of the response variables that are explained by the input variables is shown above each component by its *R*^2^ value.

Both the invertebrate community model (Fig 2; Table 3) and the functional richness model (Fig 3; Table 3) showed a strong link between the biotic metric and decomposition. Increased invertebrate community richness (i.e., higher principal coordinate values) was associated with increased decomposition rates. In this model, 32.2% of the variation in wetland decomposition was explained by the invertebrate assemblage and this was the strongest link in the model. In the functional richness model, 26.3% of the variation in decomposition was explained, with higher levels of decomposition found where there was lower functional richness.

The functional evenness model (Fig 4; Table 3), despite having satisfactory indices of fit, performed poorly in explaining the variance of measured variables relative to the other single biotic metric models (AIC = 88.294). Functional evenness was again associated with increased nutrients and macrophyte richness, with 23.5% of the total variation explained by the model. In contrast to the other two models, nutrients had a negative effect on functional evenness, with higher principal component scores associated with reduced functional evenness. The effect of macrophyte richness was also reversed, with higher levels of functional evenness associated with scores corresponding to *Vallisneria americana* Michx., and lower levels of evenness found with macrophyte species associated with plant diversity. This model revealed links between *Nutrients* and chlorophyll-*a* concentration, with increased nutrients associated with increased production, and between nutrients and decomposition, where lower principal component scores were associated with higher rates of decomposition.

**Fig 4.**
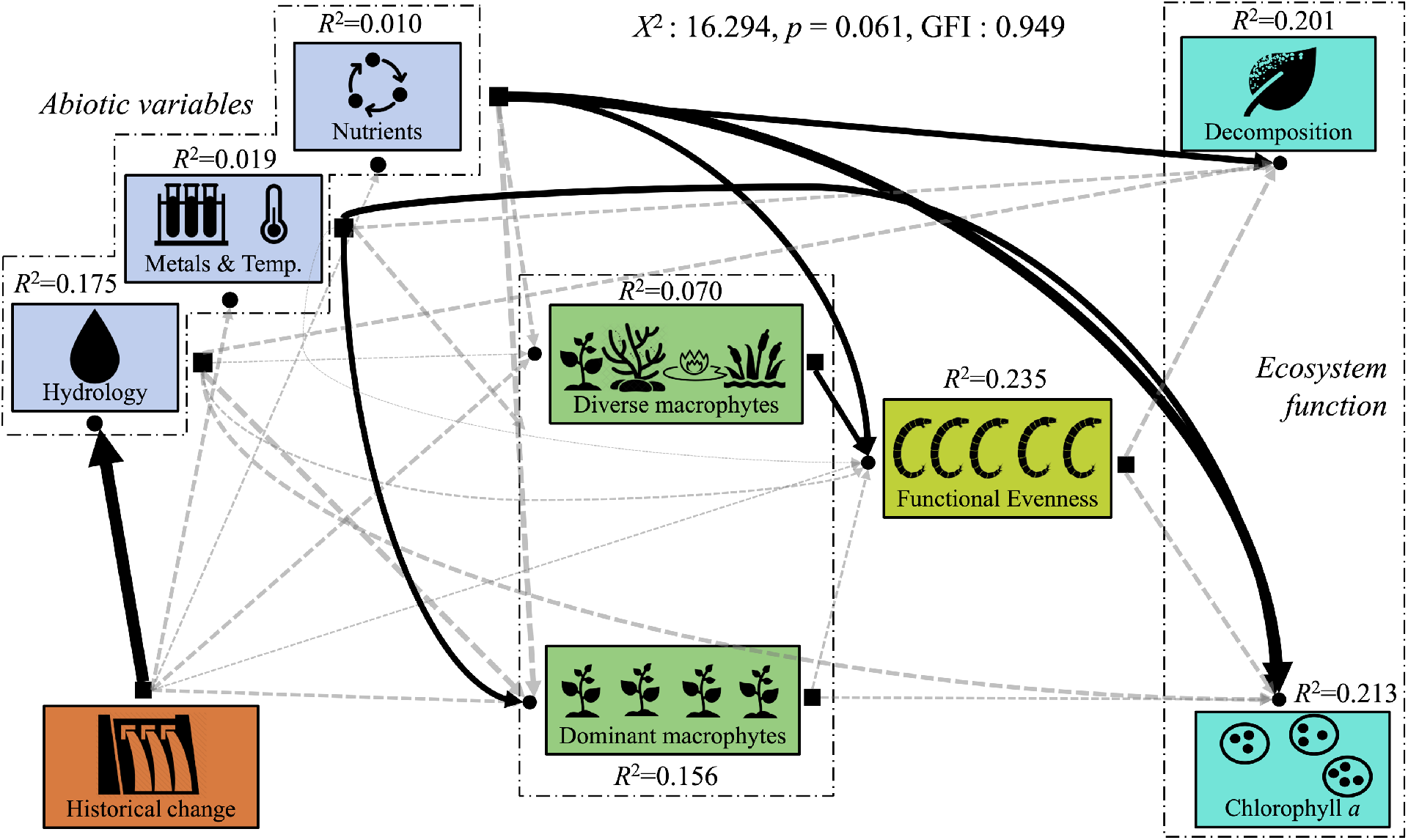
Structural equation model examining the links among the physical habitat, functional evenness, and ecosystem function. Black lines represent significant pathways at *p* < 0.05, while statistically non-significant pathways that were included in the model are depicted as dotted, gray lines. The weight of the line represents the strength of the relationship (standardized regression weights, *r*), with thicker lines having a larger effect on explaining the variation of the response variable. The variation of the response variables that are explained by the input variables is shown above each component by its *R*^2^ value.

**Fig 5.**
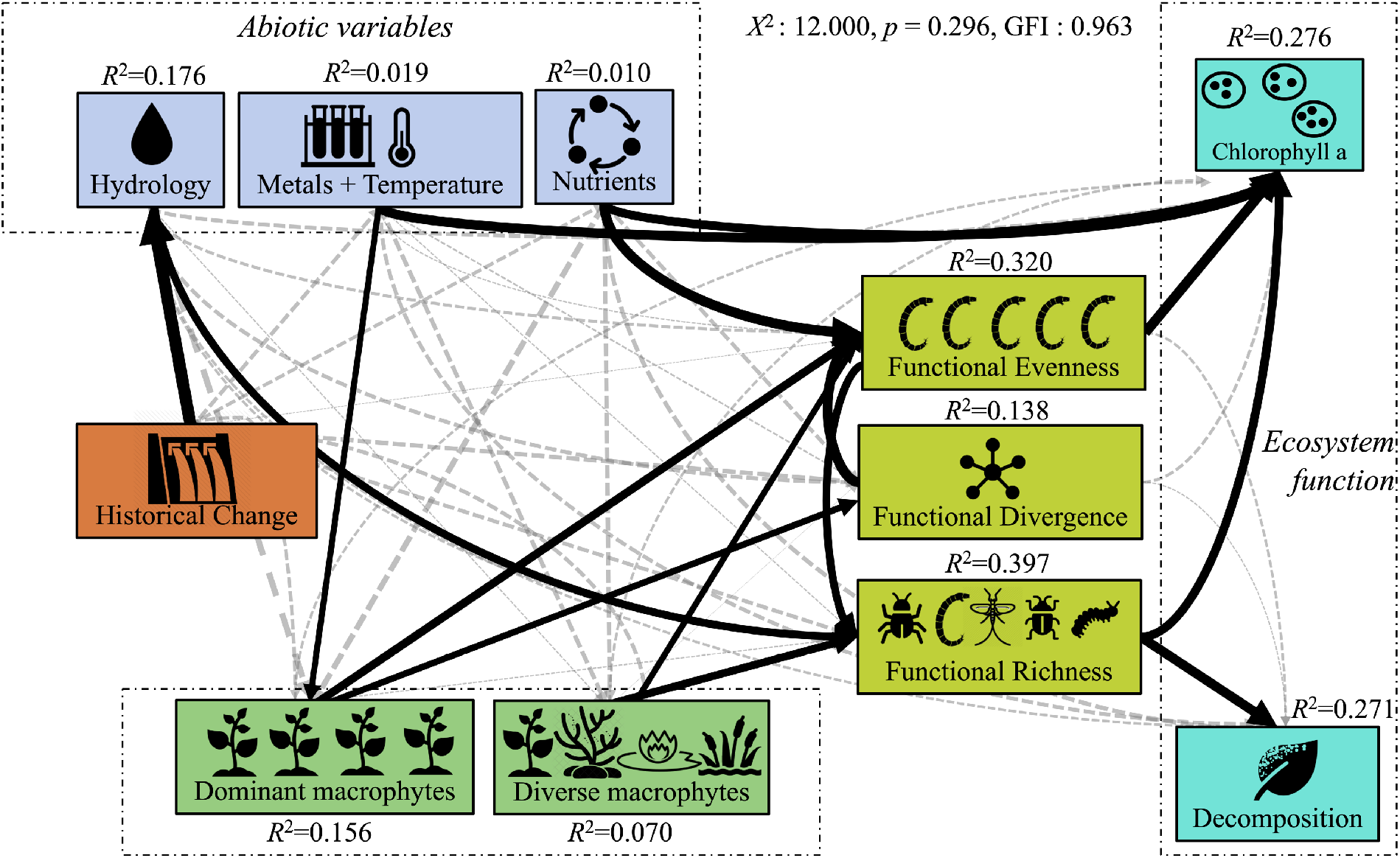
Structural equation model assessing the role of different aspects of invertebrate functional diversity in floodplain wetland ecosystem dynamics. Significant paths are shown as black, solid lines, with line weights corresponding to standardized regression weights (*r*; see Table 3.3 for estimates of the significant pathways). *R*^2^ values indicate the percent variation in a dataset that is explained by the correlative variables.

The fourth SEM accounted for all three aspects of functional diversity (richness, evenness and divergence), in a single model, allowing for direct comparisons of how traits are linked to ecosystem properties and functions (Fig 5; Table 3). All three metrics were associated with each other; although the direction of the pathway does not matter, divergence and evenness were negatively associated, as were evenness and richness. Macrophyte community dynamics had an impact on all three metrics, with sites associated with higher macrophyte diversity having lower evenness and higher richness, while sites being dominated by habitat-forming species were associated with lower evenness and divergence. Decomposition rate was not affected by any variable in the model except functional richness (accounting for 27.1% of the variation), with higher decomposition rates found at lower levels of functional richness. Primary production (*R^2^* = 0.28), estimated from chlorophyll-*a* levels as a proxy for standing biomass levels, was driven by functional evenness (i.e., evenness was positively associated with chlorophyll-*a* content), functional richness (positive association), nutrients (positive association), metals (negative association), and temperature (positive association).

### Taxa and trait indicators: TITAN2

The SEM analyses found linkages between PC1 (*Nutrients*) and decomposition rates for both invertebrate community composition (i.e. taxa) and traits. Extending this analysis, Threshold Indicator Taxa Analysis (TITAN2; [53]) was used to identify pure and reliable taxon (Fig 6) or trait (Fig 7) responses to change along the previously established gradients of abiotic and functional change. For taxa indicators, 10.67% (10 negative and 1 positive responders) were significant indicators of change in decomposition rate, compared with 12.62% (all positive responders) along PC1 (*Nutrients*) (Fig 6). Three taxa, *Hyalella, Oecetis* and *Paratanytarsus* were significant indicators for both gradients and were both positive indicators (more prevalent along the increasing gradient) for nutrients and negative responders to decomposition rate (Fig 6).

**Fig 6.**
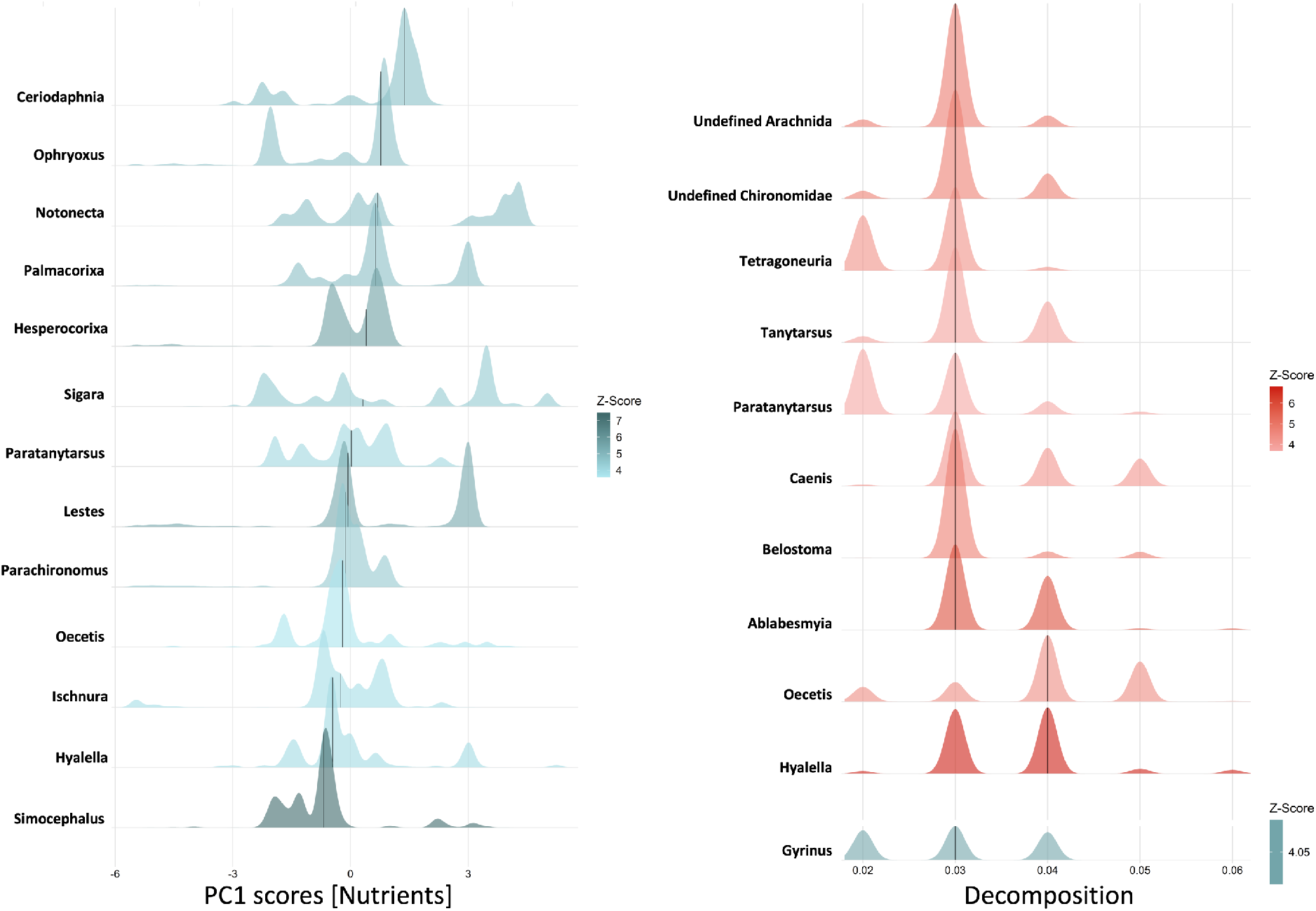
Individual response plots from Threshold Indicator Taxa ANalysis (TITAN2) comparing the response taxa modalities to changes in environmental gradients, represented by the PC axis most aligned with nutrients and ecosystem function, represented by decomposition. Taxa that responded positively to the gradient are shown in red, while negative responders are shown with blue. Taxa change points (across 999 bootstrapped replicates) are visualized as a probability density function with colour intensity scaled according to the magnitude of the response (i.e. its standardized *z*-score).

**Fig 7.**
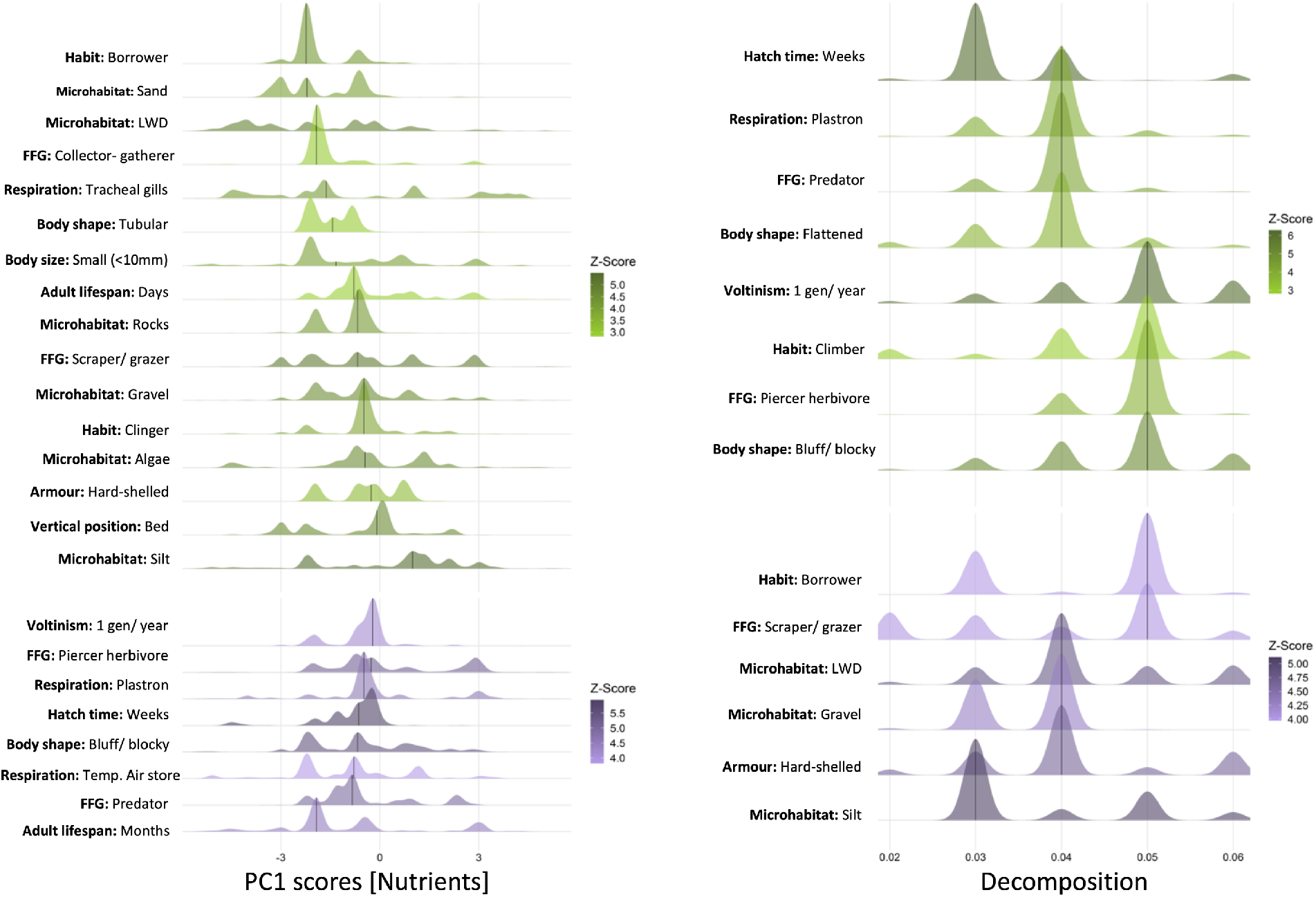
Individual response plots from Threshold Indicator Taxa ANalysis (TITAN2) comparing the response trait modalities to changes in environmental gradients, represented by the PC axis most aligned with nutrients and ecosystem function, represented by decomposition. Trait modalities that responded positively to the gradient are shown in green, while negative responders are shown with purple. Trait change points (across 999 bootstrapped replicates) are visualized as probability density functions with colour intensity scaled according to the magnitude of the response (i.e. its standardized *z*-score).

Trait modalities were better indicators of change along abiotic and functional gradients, with more pure and reliable indicators (20.90% of trait states for PC1 (*Nutrients*), consisting of 8 negative and 6 positive responders, and 35.82% for decomposition with 16 negative and 8 positive responders; Fig 7). Again, for traits, several modalities were found to be inverse indicators of PC1 (*Nutrients*) and decomposition rate. Trait states that were found to be significant positive indicators of nutrient change and negative indicators for changes along decomposition gradients were: respiration-plastron; hatch time-weeks; body shape-bluff/ blocky; Voltinism-1 generation per year; feeding mode-predator. Modalities that were significant indicators for both nutrient concentrations (negative) and decomposition rate (positive) were: microhabitat-silt; microhabitat-large woody debris; microhabitat-gravel; feeding mode-scraper/ grazer; armour-hard-shelled; and habit-burrower.

## Discussion

### The influence of disturbance on floodplain ecosystems

Flood pulses are habitat-shaping forces within floodplain wetlands, creating oxbows, swales, levees and backwaters in their erosion and deposition of sediment, and influencing productivity by replenishing adjacent wetlands with nutrients from the main river channel [18,55]. Local habitat diversity of wetland patches is thus driven by flood pulse dynamics and any alteration can influence the assemblage of organisms that live there. Using structural equation modeling, this study provides evidence that changes to flood pulse events in the Wolastoq | Saint John River, as seen from changes in wetland extent from 1951-2014, have influenced local site hydrology, leading to increased inundation during flood events, elevated temperature variation, and increased organic carbon content of the sediment. Hydrology variables were linked to changes in the functional richness of the aquatic invertebrate community, suggesting that changes to local wetland shorelines can indirectly increase functional richness.

How, exactly, could habitat change increase functional diversity? We see two potential mechanistic explanations. First, as wet meadow area shrinks, terrestrial vegetation gives way to macrophyte beds and open water (e.g. [56]), opening up habitat for more aquatic invertebrate species. The frequency and magnitude of large floods has increased over the last century, along with the likelihood that ice jam events have been altered due to increasing structures within the watershed [57]. This has resulted in increased disturbance to habitat patches within the floodplain, potentially shifting invertebrate community dynamics from being competition-to disturbance-driven [sensu 58,59]. As variability, and thus disturbance, increases along with available habitat, more species rapidly colonize vacated niche space.

Floodplain wetlands of the Platte River show similar patterns, where hydrologic regimes were key factors in shaping macroinvertebrate community structure; however, in this system hydrology gradients were linked to community dynamics of vertebrate predators, like fish [60]. Indeed, the increase in flooded aquatic wetland area also opens up habitat space for fish, which is a second mechanism that could explain how habitat change could influence functional diversity, as predators can structure invertebrate communities in a variety of other ecosystems (e.g. [61,62]. Our study did not include vertebrate predators, as it was assumed that their impact did not vary significantly across the study area because all sites were open and connected. This assumption, however, cannot be confirmed, thus predator presence may be a key component of unexplained variation within the ecosystem. An increase in functional richness supports communities with multiple avoidance strategies to limit predation, as well as providing functional redundancy so that ecosystem functions can be maintained even if one group is susceptible to external disturbances.

### Macrophytes as facilitators of invertebrate community structure

Macrophytes influence invertebrate community dynamics in several ways. They provide shelter from predatory species, like fish and birds, form numerous microhabitats within their assemblages [63], increase local oxygen levels (particularly important in wetlands, where minimal flow contributes to stagnation; although the concentration and diel patterns depend in part on the species composition of the macrophyte beds [64]), contribute to nutrient retention through increased deposition of particulate organic matter [65], and are a vital food source for many species [66]. Higher macrophyte richness increases the niche diversity of a wetland site, where species’ microhabitat preferences can range from burrowing themselves in sediment, swimming and sheltering among the complex structures of species such as *Ceratophyllum demersum* L. [66], burrowing into the inside of plant stems (e.g. larval *Euhrychiopsis* Dietz 1896, the milfoil weevil, mines into the stems of *Myriophyllum* [67]), and exploiting the floating leaves of macrophyte species (e.g. *B. schreberi*, commonly known as the water shield).

Macrophytes can even link terrestrial and aquatic communities, as evidenced by the finding of two spider genera (*Philodromus* and *Tetragnatha*), which we noticed during field collection using *Persicaria* leaves as a home base for predation of aquatic invertebrates (N. Rideout, personal observation).

In this study, macrophyte community dynamics were an important driver of the local invertebrate community, influencing both its taxonomic composition and functional diversity. Functional richness increased with macroinvertebrate diversity, tied strongly to species that were associated with highly complex macrophyte beds. The GLM complex has species rich plant communities; there were a total of 37 submerged and 15 emergent macrophyte species found in the study area (note, several species ended up being counted in both emergent and submergent surveys because of variability of individual emergence within species). The link between higher macrophyte diversity and invertebrate functional richness supported the premise that macrophytes provide more niches through increased species richness, as well as suggesting that highly diverse sites are more stable in terms of their ability to continue to provide vital ecosystem functions in the face of disturbances.

Despite their increased functional richness, however, the communities that were associated with greater macrophyte diversity also tended to have lower functional evenness within the invertebrate community. More taxa with a given trait modality were present in these macrophyte dominated sites, in contrast to more barren, exposed sites, where a variety of traits may be necessary to exploit the few niche spaces available. In these exposed sites, reduced habitat availability increases disturbance (e.g. via reduced physical attenuation of wave action) resulting in habitats where only a few species with select traits can thrive, contrasting to protective macrophyte beds where many species that are scrapers/grazers, climbers, and plant miners can coexist. Indeed, when examining abundance and richness of macroinvertebrate taxa across different macrophyte communities, Walker, Wijnhoven & van der Velde [63] found that when macrophyte complexity was high (e.g., *Elodea* and *Ceratophyllum* beds), total biomass and abundance of individuals was high but relatively few taxa dominated the samples, thus resulting in low evenness. Open water sites, i.e. those with no macrophytes, had the lowest richness and abundance as few invertebrate taxa are specialized in pelagic feeding, except for zooplankton, which are heavily impacted in these habitats by predators [63].

### Taxon richness and wetland ecosystem health

Although shredder taxa were expected to respond positively along decomposition gradients, the only positively responding indicator taxa was a predator (*Gyrinus*), likely because they are attracted to the detritivorous invertebrates for prey items. Results from TITAN2 indicate that decomposition is dominated by several species that replace each other as they compete for the food source, suggesting functional redundancy within the shredder functional group. *Hyalella* had the highest range along the decomposition gradient and in preliminary sampling for morphology and abundance the previous year was found to dominate the samples in which it was present (N. Rideout, unpublished data). Taken in accordance with SEM results, which show that functional richness decreased with increasing decomposition, this suggests that several prevalent species are responsible for the majority of invertebrate-mediated decomposition, outcompeting other functional groups.

In freshwater ecosystems, the majority of decomposition studies have been carried out in the main channels of rivers and small streams, where leaf litter input is rapid and substantial in the autumn, opening up niche space for shredder diversity. In contrast, litter inputs on floodplains are more gradual, with leaf litter moving into the aquatic wetland from the floodplain forest (e.g., the silver maple floodplain forest of the GLM complex) with the flood pulse, settling on the soft bottom of the wetlands [68]. In temperate systems, there may be a significant time lag between leaf litter fall and its eventual decomposition, with settled particulate organic matter creating standing crops that can be as much as 6-7 years old [68]. The nature of this input means that leaf litter is not necessarily a limited resource in wetlands, leading to several species specializing on it as a food source and, thus, dominating the community. Moreover, through the amassing of trait data, it was noted that there were many more taxa that relied on algae or fresh macrophytes as a food source than shredding detritus (e.g. Hydroptilidae (i.e. microcaddisflies or purse-case caddisflies), mayfly families such as Baetidae and Heptageniidae, and larval Elmidae (riffle beetles)). Tiegs *et al.*, [69] also noted that leaf-shredding invertebrate taxa were rare or absent in their study along the Tagliamento floodplain, and Cuffney [68] noted that floodplain communities along the Ogeechee River were rich in collector-gatherers and collector-filterers, but poor in shredders. In lotic ecosystems, macroinvertebrates are the dominant source of leaf litter breakdown with shredders shown to contribute over half of the overall leaf mass loss [69]. In floodplains, however, fungi are much more important, contributing about 49% of the decomposition function, in contrast to an estimated 41% in rivers and streams [70]. In this study, it was notable that overall richness of the invertebrate assemblage was not strongly associated with decomposition in wetlands, contrasting the situation observed in streams and rivers, and that the shredder trait modality was not a significant indicator of decomposition in TITAN2 analysis. Scrapers and grazers do positively respond to increased decomposition, supporting the theory that floodplain decomposition is fungal-dominated and so increased fungal biomass on leaf litter can attract a wider range of species which can exploit them as a food source.

While decomposition was associated with negatively responding taxa, nutrients were only associated with positive responders. Notably, multiple predator taxa acted as indicators of increased nutrients. Healthy ecosystems can support more predators (e.g. [71]), suggesting that in the Grand Lake Meadows and Portobello Creek wetlands, increases in nutrients like nitrogen and phosphorus were associated with increased ecosystem health. Floodplains tend to be phosphorus- and nitrogen-limited and are dependent on flood pulses to replenish these nutrients into the wetlands from the main channel, whence they increase algae productivity and support macrophyte communities [72]. From SEM analysis, we know that higher levels of nutrients are positively associated with functional richness, along with increased invertebrate community diversity. This suggests that wetland sites with higher nutrients support higher functional redundancy and, despite having lower local levels of decomposition, are more stable in their ability to maintain functional roles in the face of disturbance.

The inverse relationship between nutrients and decomposition is supported by the observation that multiple taxa and trait modalities were inverse indicators of decomposition and nutrients. Most notable in this inverse relationship were a) predators, which positively respond to increased nutrients, b) microhabitat modalities associated with burrowing, which are positive indicators of decomposition, and c) scrapers and grazers, which are positive indicators of decomposition. Again, this indicates that in wetlands, healthy and productive sites are those with high nutrients, algal production, and dense macrophyte beds, and that these sites are those associated with high invertebrate functional richness. Primary productivity, as assessed by proxy through chlorophyll-*a* content of periphyton on planted tiles, increased with increasing nutrients, as well as with functional evenness. In functionally-even communities, resources are used efficiently with no single trait modality dominating the community, implying, for example, that no taxa rely heavily on grazing algae, therefore allowing for increased algal standing biomass. It is worth remembering, however, that chlorophyll-*a* is a static estimate of periphyton standing crop and does not necessarily reflect instantaneous primary productivity at a site; for example, a site could be highly productive, with high algal turnover due to invertebrate grazing and so have a relatively low chlorophyll-*a* value.

### Conclusion: trait metrics for biomonitoring

While the SEM model examining the linkage in the ecosystem with invertebrate community composition had the best fit when compared to those incorporating multidimensional trait metrics, the overall information provided by trait data better explained ecological inter-relationships when compared to using taxonomic information alone. Coupling this approach with DNA metabarcoding reveals a more complete picture of invertebrate community composition than provided by traditional microscope-based observations (e.g. [24]). This is significant, as incomplete observational coverage has been a limitation for traits work in the past. One drawback with DNA-based community data is that information is in the form of presence/ absence, so despite knowing how many taxa hold a certain trait in a community, we are limited in that we do not know the dominance of that trait in terms of the actual abundance or biomass at each site [73,74].

Traits simplify communities into generic groupings, which facilitate cross-study comparisons [75]. By partitioning traits into those that affect ecosystem function and those that respond to abiotic changes, we can start to make inferences about specific links between ecosystem components [14]. This was evident when examining the output from TITAN2 analysis; trait data revealed critical insights into how individual taxa were responding along these gradients, revealing information about poorly studied ecosystem response pathways in wetland ecosystems, particularly the role of invertebrate taxa in decomposition processes.

By measuring trait distribution in multidimensional space, we can examine niche complementarity and resource distribution, a proposed mechanism behind the link between diversity and ecosystem function [75]. We can think of niche space as the volume of the convex hull created by each site’s calculation of functional diversity from the PCoA; aspects about a community’s multidimensional trait space can be compared across the ecosystem at the same scale. Higher functional richness results in a greater volume of niche space and thus a more complete use of resources, whereas the distribution of traits in the niche space (i.e. its evenness) determines the heterogeneity of resource use [76]. Functional divergence indicates how much a community has maximized its total variation in the niche space though the distribution of taxon abundance and is thus an indication of the efficiency of resource use [76]. Functional divergence indicates competition [76] and can therefore be thought of as an indicator of functional redundancy.

Healthy ecosystems—those which maximize ecosystem function and resilience—can achieve stability through balancing efficiency and completeness of resource use with functional redundancy. By providing information about the efficiency of resource use within an ecosystem and the amount of competition present, functional divergence is a potent indicator of ecosystem health; in our system, however, without abundance information, our measure of functional divergence lacks power [32]. Yet, even in the absence of functional divergence information, measures of high functional richness and evenness can indicate healthy ecosystems. Communities that show high functional richness can be resilient under environmental fluctuations, since the taxa with traits necessary to take advantage of new conditions are more likely to be present [76]. Comparatively, functionally-even communities are efficiently utilizing the entire range of the ecosystem’s resources, reducing the opportunity for foreign invaders to occupy niche space [76]. In disturbance-dominated systems, such as river floodplains, the ability to respond to, and to buffer against, environmental fluctuations enables communities to support and sustain the vital ecosystem functions provided to society by floodplains and their associated wetlands.

## Acknowledgments

We thank members of CRI and ECCC for valuable discussions and feedback in developing our study and manuscript. Additionally, we owe many thanks to K. Heard for her invaluable help and expertise in the field, and to S. Connor, and R. Anema for assisting with field and lab work. We also thank members of the New Brunswick Department of Energy and Resource Development for access to historical aerial imagery and to S. Stefani, Z. O’Malley, C. Brooks, and J. Ogilvie for their parts in the associated data gathering and analysis.

## Data availability statement

Raw sequence data has been deposited to the National Centre for Biotechnology Information (NCBI) Short Read Archive (SRA) under BioProjectID PRJNA640405. The bioinformatic pipeline used to process COI metabarcodes is available from GitHub at https://github.com/Hajibabaei-Lab/SCVUC_COI_metabarcode_pipeline. The COI Classifier used to make taxonomic assignments is also available from GitHub at https://github.com/terrimporter/CO1Classifier. All other data will be made available on an online repository (e.g. Dryad) upon paper acceptance.

## Supporting information

**S1 Table. Pathways from structural equation models examining the relationship between individual ecosystem components.** All pathways are listed, including statistically non-significant ones, with their significance indicated by *p* and their standardized regression weight indicated by *r*. In total we tested four structural equation models.

